# Redesigned Reporter Gene for Improved Proton Exchange-based Molecular MRI Contrast

**DOI:** 10.1101/2020.04.21.053157

**Authors:** Or Perlman, Hirotaka Ito, Assaf A. Gilad, Michael T. McMahon, E. Antonio Chiocca, Hiroshi Nakashima, Christian T. Farrar

**Author notes:** Correspondence to: Christian T. Farrar, Athinoula A. Martinos Center for Biomedical Imaging, Department of Radiology, Massachusetts General Hospital, 149 13th Street, Suite 2301, Charlestown, MA, 02129, USA.

## Abstract

Reporter gene imaging allows for non-invasive monitoring of molecular processes in living cells, providing insights on the mechanisms underlying pathology and therapy. A lysine-rich protein (LRP) chemical exchange saturation transfer (CEST) MRI reporter gene has previously been developed and used to image tumor cells, cardiac viral gene transfer, and oncolytic virotherapy. However, the highly repetitive nature of the LRP reporter gene sequence leads to DNA recombination events and the expression of a range of truncated LRP protein fragments, thereby greatly limiting the CEST sensitivity. Here we report the use of a redesigned LRP reporter (rdLRP), aimed to provide excellent stability and CEST sensitivity. The rdLRP contains no DNA repeats or GC rich regions and 30% less positively charged amino-acids. RT-PCR of cell lysates transfected with rdLRP demonstrated a stable reporter gene with a single distinct band corresponding to full-length DNA. A distinct increase in CEST-MRI contrast was obtained in cell lysates of rdLRP transfected cells and in *in vivo* LRP expressing mouse brain tumors (p=0.0275, n=10).

## Introduction

Magnetic resonance (MR) reporter genes can non-invasively monitor transgene expression *in vivo* at high resolution with unlimited tissue penetration^1^. These genes can be utilized for the evaluation of cell and gene-based therapy, as investigated for cancer, neural degeneration, and myocardial infarction^2^. Specific applications include tracking of dividing/differentiating stem cells^3^, imaging of dendritic cells^4^, monitoring of viral therapeutics^5^, and detection of micro-environment changes^6^.

The adaptability of MRI and the different contrast mechanism that can be used, fostered creativity among scientists that allowed the development of numerous genetically encoded reporters^7, 8^. In a pioneering work in the field, Louie et. al., used beta-galactosidase to catalyze an MRI specific probe^9^. This study set the stage for a variety of genetically encoded reporters based on T_1_ relaxation. Bartelle, Turnbull and their colleagues have taken a creative approach to express a multi-component reporter specifically in endothelial cells^10^. Another strategy relied on overexpression of the organic anion transporting protein (Oatp1a1). These transporters facilitate the uptake of Gd(3+)-DTPA^11, 12^. A different approach was to develop genetically encoded reporters based on T_2_ relaxation. By using iron transporters^13–15^ or iron storage proteins^16–18^, scientists sought to modulate the intracellular T_2_ relaxivity. An alternative approach was to use hyperpolarized xenon as a substrate for enhancing the MRI contrast of a genetically encoded reporter^19^. Nevertheless, all these innovative approaches relied on administration of a substrate or a secondary compound that is not accessible to all tissues and requires repetitive injections for longitudinal imaging.

Chemical exchange saturation transfer (CEST) is an alternative and particularly useful contrast mechanism for MR reporters^20–25^. In a typical CEST-MRI experiment, compounds containing exchangeable protons are selectively saturated using selective radio-frequency (RF) irradiation and undergo chemical exchange with the bulk water protons. A signal associated with the compound concentration and its labile proton exchange rate can then be detected using the water signal^26–28^. CEST MRI reporter genes offer a few attractive advantages: (i) The artificially engineered CEST proteins are non-metallic and bio-compatible. (ii) The imaging procedure does not require the injection of a concomitant labeling agent. (iii) The reporter contrast can be switched on/off by the use of a selective MR RF saturation pulse, allowing the acquisition of additional and uncontaminated anatomical/functional MRI information at the same imaging session. Of particular promise was the development of the lysine-rich protein (LRP) reporter gene^23^, with demonstrated applicability for imaging cells in tumors^29^, oncolytic virus delivery and spread^5^, and cardiac viral vector expression^30^. However, thus far, the gap between the promise of MR reporters and actual clinical use has not been bridged due to the low sensitivity and specificity of the detection. In particular, the LRP reporter gene in its current form is unstable, as the highly repetitive DNA sequence encoding the reporter leads to DNA recombination events and hence the expression of a range of truncated protein fragments with fewer lysine amide protons, resulting in decreased and limited CEST sensitivity^5, 29^.

Here, we report the redesign of an improved, stable, and robust LRP-based CEST-MRI reporter gene (rdLRP), using an RHGP amino acid motif that consisted of arginine, histidine, glycine, and proline amino acids separated by different numbers of lysines. The reporter stability and contrast enhancement ability were evaluated in cell lysates from human embryonic kidney (HEK293T) and murine glioma (GL261N4) cell lines as well as from *in vivo* LRP expressing brain tumors in mice.

## Results

### CEST-MRI of various polypeptides

A number of different lysine polypeptides with different amino acid motifs separating lysine residues were screened to find a peptide that disrupts the repetitive lysine sequences and has optimal CEST contrast. Of these, the CEST contrast of a lysine polypeptide containing an arginine, histidine, glycine, and proline (RHGP) amino acid motif stood out as this displayed virtually the same CEST contrast as poly-L-lysine (PLL) for the amide exchangeable protons at 3.6 ppm chemical shift (Fig. 1). In addition, a strong CEST contrast was observed at the amine chemical shift of 1.8 ppm attributed to the guanidyl amines of the arginine residues (Fig. 1a). Based on the contrast advantages described above, the rdLRP was engineered using RHGP units with different lysine spacers and a total of 100 amino acids (Fig. 2a).

**Fig. 1.**
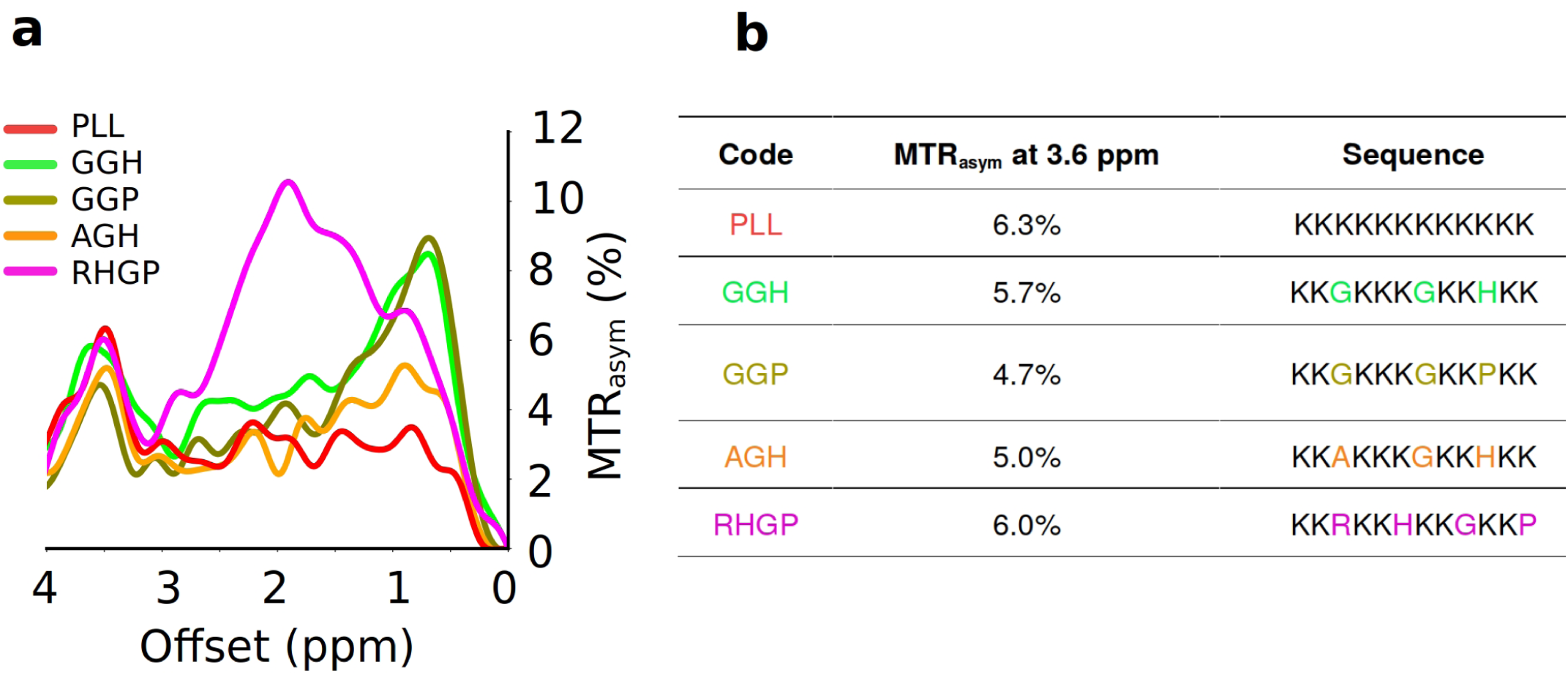
Screening various polypeptides in search of an optimal LRP-based reporter gene backbone. **a**. CEST contrast observed as a function of saturation frequency offset for a variety of polypeptides with different amino acid motifs. **b**. Polypeptide sequences and their respective CEST contrasts at the amide exchangeable proton chemical shift of 3.6 ppm. The amide proton CEST contrast for the RHGP polypeptide (containing the arginine, histidine, glycine and proline amino acid motif) is similar in magnitude to that of PLL. A large CEST contrast is also observed at 2 ppm for the amine exchangeable proton of the RHGP polypeptide. A: Alanine, PLL: poly-L-lysine.

**Fig. 2.**
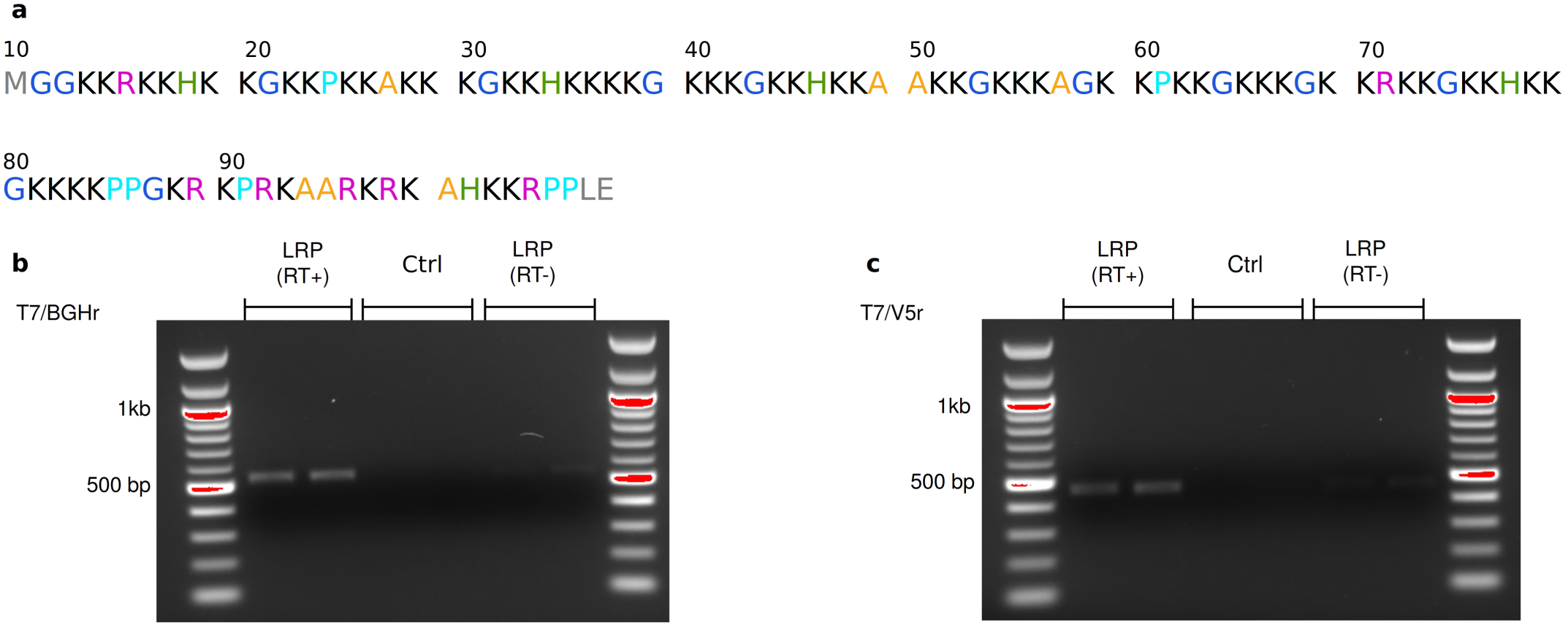
RNA characterization of the rdLRP reporter. **a**. Amino acid sequence of the rdLRP consisting of RHGP units with different lysine spacers with a total of 100 amino acid residues. **b-c**. RT-PCR of total mRNA from HEK293T cell lysates transfected with rdLRP, evaluating the BGH reverse tag (**b**) and the v5 epitope tag (**c**) sites. A single well-defined band is observed from cell lysates transfected with rdLRP, but not from control cell lysates. The base-pair lengths are consistent with expression of full-length rdLRP mRNA (**b**, 557 bp; **c**, 491 bp).

### Stability analysis of the rdLRP reporter

The rdLRP contains no DNA repeats or GC rich regions resulting in a significantly more stable protein with 30% less positively charged amino acids compared to the original LRP (see Extended Data Fig. 1). The improved stability is evident in RT-PCR of total mRNA from cell lysates transfected with rdLRP, where only a single well-defined DNA band is observed at the base pair length expected for full length rdLRP (Fig. 2b-c). In contrast, a ladder profile of bands of different base-pair lengths was observed previously in RT-PCR of cell lysates transfected with the original LRP reporter gene (see Fig. 2f in^23^ and Fig. E2 in^5^).

### *In vitro* CEST-MRI of rdLRP transfected cell lysates

A distinct increase in the CEST magnetization transfer ratio asymmetry (MTR_*asym*_) was observed for both human embryonic kidney (HEK293T) cells and murine glioma (GL261N4)^31^ cells transfected with the rdLRP relative to control, non transfected cells (Fig. 3). The two main amplitude peaks in the MTR_*asym*_ of the cell lysates were at the amine (∼1.8 ppm) and amide (∼3.6 ppm) exchangeable proton frequencies, in agreement with the RHGP peptide CEST analysis (Fig. 1).

**Fig. 3.**
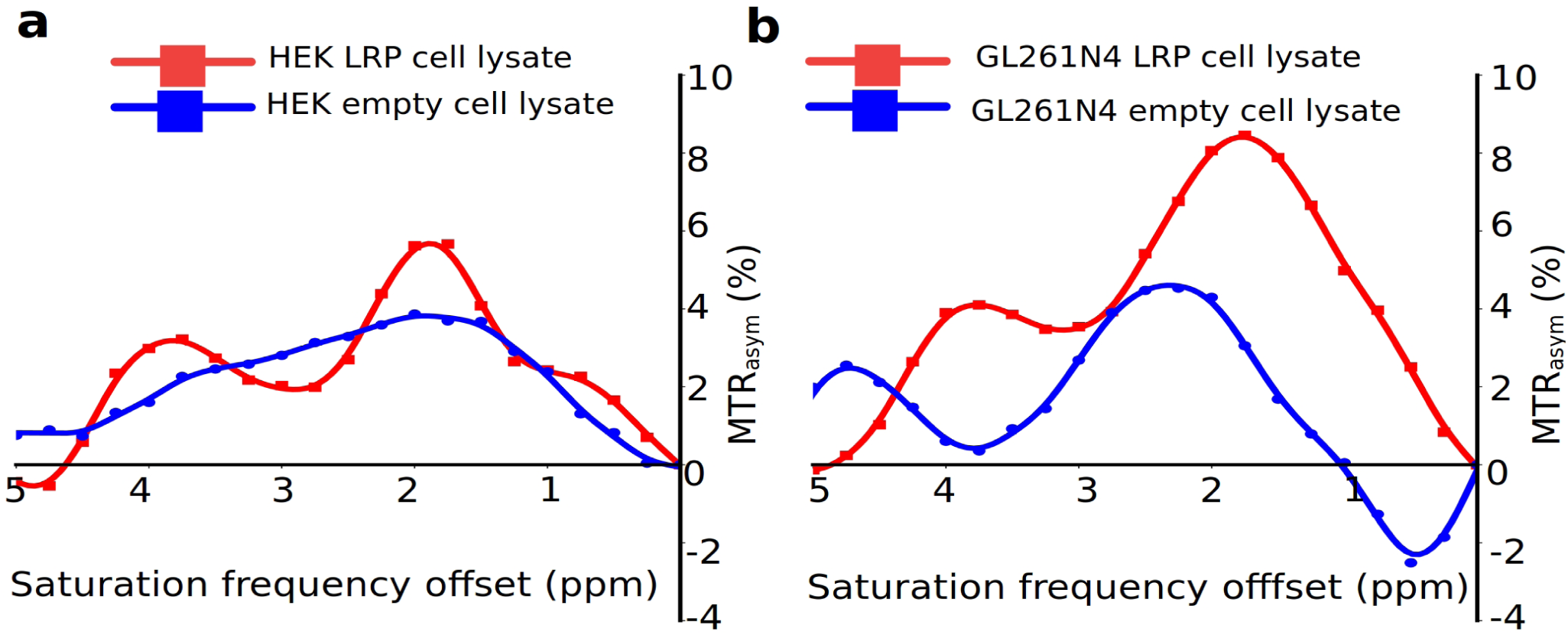
*In vitro* CEST-MRI contrast evaluation of the rdLRP reporter. **a-b**. MTR_*asym*_ of control (blue trace) and rdLRP (red trace) in transfected HEK293T (**a**) and GL261N4 (**b**) cell lysates acquired at 4.7T. Increased CEST contrast is observed at ∼1.8 ppm corresponding to the guanidyl amine exchangeable protons of arginine amino acid residues and at ∼3.6 ppm corresponding to the amide exchangeable protons of lysine amino acid residues.

### *In vivo* imaging of rdLRP expressing mice brain tumors

To verify that the contrast enhancement ability of the rdLRP reporter is retained for *in vivo* settings and to demonstrate the reporter effectiveness for a molecular-genetic imaging scenario, a preclinical tumor monitoring experiment was conducted. The rdLRP was expressed in a murine glioma GL261N4 cell line and transplanted into mouse brains. *In vivo* CEST-MR images were acquired at 7T and the amide-proton based contrast was analyzed (Fig. 4). As the molecular environment of the living brain/tumor has a more diverse multi-compound content compared to *in vitro* cells, the extraction of the CEST amide proton amplitude was performed using a Lorentzian fitting model (see the Methods section), aimed to isolate the signal of interest from the confounding contribution stemming from lipid/semi-solid macro-molecules and aliphatic protons (rNOE). A significant increase in the tumor amide-CEST contrast was obtained for LRP expressing tumors (5.88±1.44%, mean±STD) compared to control (4.34±1.28%, mean±STD), as shown in Fig. 4f (two-tailed t-test, p=0.0275, n=10 imaging data points, t=2.399, df=18, difference between means: 1.542±0.643% (mean±SEM)). These findings demonstrate that the rdLRP can be expressed *in vivo* in a cell-selective manner and can be used for molecular MR studies. It shall be noted, that due to evident edema (Fig. 4c-e), the protein content in the tumor region was diluted, resulting in a general decrease in the CEST signal compared to the contralateral region (Fig. 4a-b, Extended Data Fig. 2), as previously reported in other CEST tumor studies^32–34^. The edema has similarly affected the control and the rdLRP group mice, as reflected by the similar increase in T_2_ value for both groups (from 48.18±0.75 ms to 72.86±12.05 ms for the contralateral and tumor ROIs in the control mice group, respectively, and from 48.33±0.79 ms to 73.08±7.92 ms in the rdLRP mice group contralateral and tumor ROIs, respectively). The difference between the control and rdLRP mice groups T_2_ tumor values was not statistically significant (two-tailed t-test, p = 0.961, t = 0.0491, df = 18, n = 10 imaging points per group). Hence, although sharing similar conditions for the endogenous amide content and edema extent, the rdLRP mice group has presented a distinct amide CEST contrast increase, resulting from the presence of the rdLRP.

**Fig. 4.**
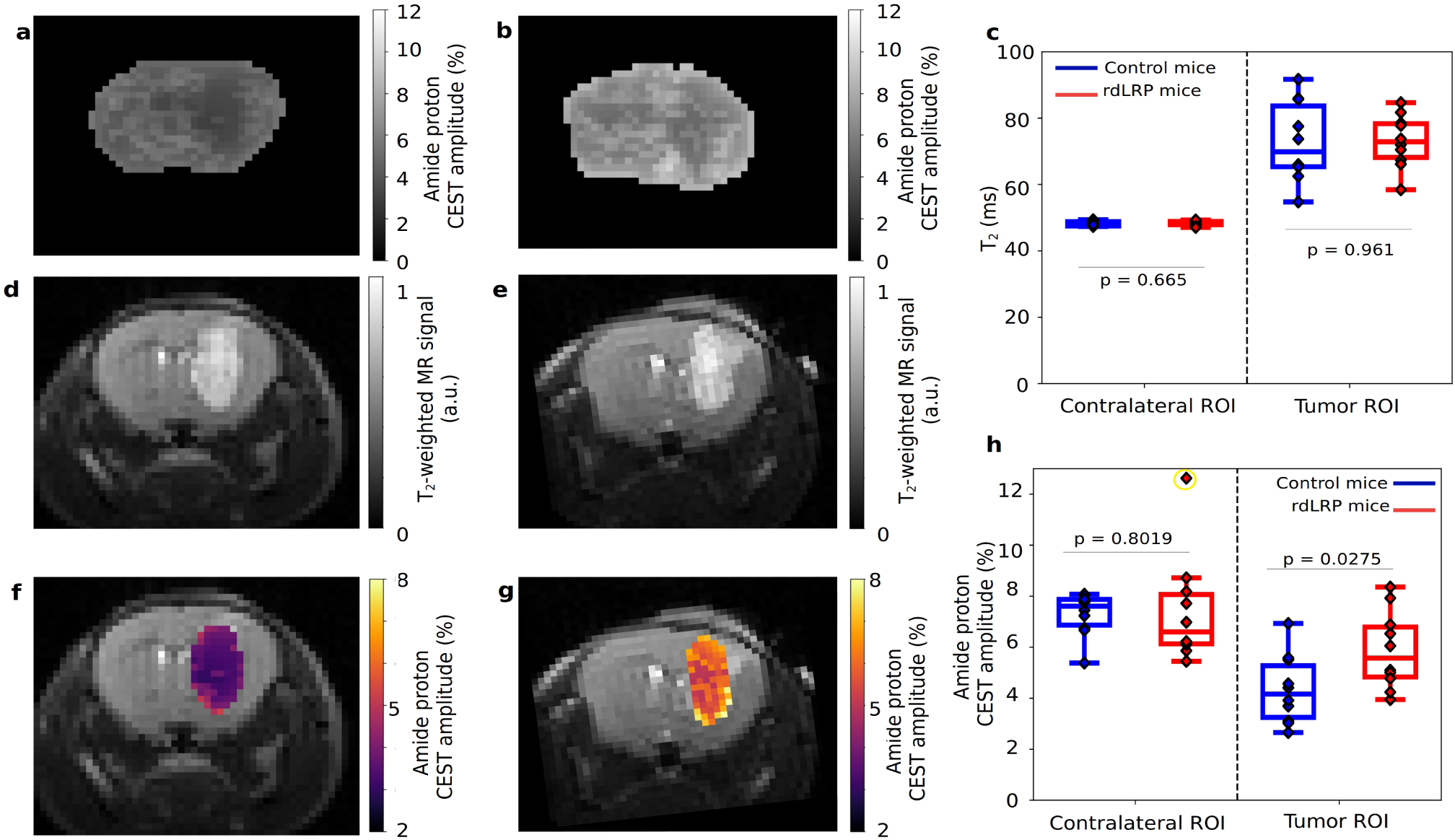
*In vivo* imaging of the rdLRP. **a-b**. Amide proton CEST amplitude maps of representative mice implanted with either GL261N4 glioma cells (**a**) or with rdLRP transfected GL261N4 cells (**b**). **c**. T_2_ relaxation time analysis for the two imaged groups, demonstrating the evident edema at the tumor region. Notably, no statistically significant differences were calculated at the tumor region between the rdLRP and control group mice (two-tailed t-test, p=0.961, t=0.0491, df=18, n=10 imaging points per group). **d-e**. T_2_-weighted images for the mice imaged at **a-b**, respectively. **f-g.** The amide proton CEST contrast, obtained using a frequency selective saturation pulse (**a, b**) is overlaid atop the tumor region in the T_2_-weighted images presented in **d** and **e**, respectively. **h.** Quantitative group comparison for the amide proton CEST amplitude. Statistically significant CEST signal increase was obtained in the rdLRP group (two-tailed t-test, p=0.0275, n=10 imaging data points, t=2.399, df=18, difference between means: 1.542±0.643% (mean±SEM)). The box plots represent the median, interquartile range, Tukey style whiskers, outliers (yellow) and 27 individual data points.

## Discussion

Stability and reproducibility are two essential factors required for the successful application of a reporter gene^35^. Furthermore, to allow the correct quantification of the biological/therapeutic process of interest (e.g., the extent of viral replication and spread during oncolytic virotherapy^5^), it is essential to reliably know the exact composition of the contrast-generating substance in the reporter (e.g., the number of amide bonds of the backbone, as affected by the environment induced by the presence of lysine amino acids) and avoid contaminating effects, such as random protein truncation. However, engineering highly predictable and reliable reporters is technically challenging, as demonstrated in the gene expression imperfections of previously developed CEST-MRI reporter genes^5, 23^. Here, the LRP reporter was redesigned for overcoming its previous limitations and enhance its performance. The results have demonstrated a marked improvement in stability, as evident in RT-PCR (Fig. 2), where only full-length cDNA was observed.

The rationale of this study was to increase the LRP cellular expression and make it a more stable reporter. This will allow to use a smaller protein, half the size of the original LRP, and still ensure sufficient contrast. This is a major consideration when designing a viral gene delivery system with limited DNA payload. To that end we modified the LRP at the translational, transcriptional and protein level. For transcriptional efficacy we minimized the GC content and eliminated CpG dinucleotides content, cryptic splicing sites, negative CpG islands and TATA boxes. For improving translation efficiency, we optimized codon usage for mammalian cells. This approach was proven very efficient in the expression of another MRI reporter gene, the human PRM1, where codon optimization turned the protein from practically unexpressed to highly expressed in E. Coli^20^. The new rdLRP sequence (Extended Data Fig. 1) also encodes to less than 60% lysines, unlike previous generation that encodes close to 100% lysines and was double in length. In other words, the rdLRP uses only 35% of the number of Lysines residues per protein. Assuming a strong gene promoter is used, which is often the case, the original sequence may deplete the cellular lysine reservoirs and slow down the translation by overusing the lysine-tRNALys. Therefore, reducing the lysine content can significantly reduce the burden on the cellular protein production machinery. In addition, we eliminated mRNA secondary structures, RNA instability motifs, removed ribosomal binding sites and minimized protein refolding. We incorporated protein motifs that were suggested to increase peptide resistance to degradation and improve peptides stability^36^. Together, these manipulations, lead to a more stable LRP that is smaller but can be overexpressed in higher concentrations in cells and consequently increase the reporter detectability.

Importantly, the CEST-MRI contrast enhancement ability was fully retained compared to the original LRP reporter gene. The *in vivo* CEST contrast improvement calculated in this work (1.542±0.643%, mean±SEM) is similar to that reported in previous LRP studies, where an *in vivo* increased CEST contrast of 1.1%^5^ or less^29^ was observed. As mentioned earlier, the smaller rdLRP reporter gene contains only 35% of the total lysine content theoretically available in the original LRP reporter^23^. However, this change in the reporter gene composition is compensated by its superior stability, allowing to de facto take advantage of its full contrast enhancement potential, without any unexpected decreases in sensitivity due to random DNA/protein truncation. With a stable reporter gene on hand, the size of the reporter gene could be extended for a further gain in CEST sensitivity.

Another beneficial contrast feature of the rdLRP was demonstrated in its CEST MTR_*asym*_ (Fig. 3), where an additional and strong CEST contrast peak was observed at the amine chemical shift of 1.8 ppm, attributed to the guanidyl amines of the arginine residue. This may provide a further boost in the CEST sensitivity as well as provide a means for performing amine/amide ratiometric CEST imaging for obtaining quantitative pH measurements independent of the reporter concentration (as performed for iodine based CEST contrast agents^37, 38^). In addition, the very rapid exchange rate (1 kHz) of the amine exchangeable protons may provide a means for distinguishing the CEST reporter from endogenous proteins with slow proton chemical exchange by exploiting exchange rate specific CEST methods such as Frequency Labeled EXchange transfer (FLEX)^39^. However, the resonance frequency of amine protons is close to water. Therefore, to avoid direct water saturation, the saturation pulse power must be limited, preventing the sufficient amplification of the amine fast exchanging proton signal. A potential solution might come from implementing a multi-saturation power acquisition scheme, as performed in CEST MR-fingerprinting^40–42^, with the potential addition of CEST neural networks for separating out the contribution of each contrast-generating compound^34^.

In summary, a second generation of the LRP CEST reporter gene was engineered, characterized, and thoroughly evaluated using RT-PCR, cell lysates from two cell lines, and *in vivo* LRP-expressing tumors in mice. Significant improvement in stability and contrast enhancement ability were demonstrated. Future work is underway to optimize the protein sequence even more with emphasis on reporters that generate CEST contrast at 5 ppm where the endogenous CEST background is lower.

## Conclusion

The redesigned LRP reporter demonstrates excellent stability with distinct CEST contrast observed for both amide and guanidyl amine exchangeable protons of the redesigned reporter. The redesigned reporter gene was detectable *in vivo* in a mouse tumor model engineered to express the rdLRP.

## Acknowledgments

The work was supported by the US National Institutes of Health Grants R01-CA203873, P41-RR14075, as well as by 1R01NS110942-01A1 and P01 CA163205 (EAC and HN) and NIH R01 NS098231; R01 NS104306 and P41 EB024495 (AAG). The Brigham and Womens Small Animal Imaging Laboratory (SAIL) was funded by a G20 Grant (1G20RR031051-01) as part of the American Recovery and Reinvestment Act as a part of the construction of the Brigham and Womens MRI Research Center (BWMRC). The 7T Bruker Small Bore Animal Magnet was partially funded by an S10 Grant (1S10OD010705-01) through the National Institutes of Health. This project has received funding from the European Union^*t*^s Horizon 2020 research and innovation programme under the Marie Sklodowska-Curie grant agreement No. 836752 (OncoViroMRI).

## Author contributions

A.A.G., M.T.M., and C.T.F. conceptualized the problem. O.P., H.I., A.A.G., M.T.M., H.N., E.A.C., and C.T.F. contributed to experimental design. A.A.G. designed the reporter gene. M.T.M. performed the peptide imaging study. H.I. and C.T.F. performed and analyzed the cell lysates study. H.I. performed the mice tumor implantation. O.P., H.I., and C.T.F. performed the *in vivo* imaging. O.P. and C.T.F. analyzed the *in vivo* data. O.P., H.I., A.A.G., M.T.M, H.N., E.A.C., and C.T.F. wrote and/or substantially revised the manuscript.

## Methods

### CEST-MRI of various polypeptides

Five synthetic peptides (Genscript, Piscataway, NJ) were dissolved in PBS (Fig. 1) and their CEST Z-spectra were obtained on a 11.7T vertical-bore MRI scanner (Bruker Biospin, Ettlingen, Germany) with imaging parameters as described previously^23, 29^. Briefly, peptides were placed in 5 mm NMR tubes and the collection placed within the scanner at 37°C. A modified rapid acquisition with relaxation enhancement (RARE) sequence was implemented, with repetition time/echo time (TR/TE) = 5000/20 ms, RARE factor = 16, slice thickness = 1mm, field of view (FOV) = 1.7 × 1.7 cm, matrix size = 128 × 64, resolution = 0.17 × 0.34 mm, and a saturation pulse (amplitude = 3.6 *µ*T, duration = 3s), repeated over a frequency offset range of −5 ppm to 5 ppm with 0.1 ppm steps. The water saturation shift reference (WASSR) method^43^ was used for B_0_ corrections, using a similar protocol with TR = 15s, saturation pulse amplitude, duration, and range of 0.5 *µ*T, 0.5s, and −1 ppm to 1 ppm with 0.1 ppm steps, respectively.

### Redesigned LRP (rdLRP) reporter gene synthesis

The rdLRP was designed using the peptides in Fig. 1 as a starting point for the assembly of the gene with emphasis on amino acid motif that consisted of arginine, histidine, glycine, and proline amino acids separated by different numbers of lysines. Next, the gene was optimized by considering transcriptional efficacy (i.e., GC content, CpG dinucleotides content, cryptic splicing sites, negative CpG islands and TATA boxes) as well as translation efficiency (including: codon usage bias, GC content, mRNA secondary structure, RNA instability motif, stable free energy of mRNA and ribosomal binding sites) and protein refolding (e.g., codon usage bias, interaction of codon and anti-codon and RNA secondary structures). For that end we use dedicated algorithms from GenScript (Piscataway, NJ). A full length DNA sequence of the gene was custom synthesized by GenScript (Piscataway, NJ) and was further subcloned into expression vector.

### Stability characterization of the rdLRP reporter

The Taq polymerase-amplified PCR product of the rdLRP was inserted into the TOPO Cloning site of a pcDNA3.1/V5-His-TOPO vector (Thermo Fisher Scientific, MA). The vector was transfected to 6×105 cells of HEK293T cells using Opti-MEM (Thermo Fisher Scientific, MA) and a Lipofectamine reagent (Thermo Fisher Scientific, MA) according to the vendors protocol. RNA was extracted from the transfected cell pellets using TRIzol reagent (Thermo Fisher Scientific, MA). Purified RNA template was reverse-transcribed to cDNA by using iScript cDNA Synthesis Kit (Bio-Rad, CA) and polymerase chain reaction (PCR) was performed on T-100 Thermal Cycler (Bio-Rad, CA). Two kinds of combinations of primers (for T7 priming site/BGH Reverse priming site, and for T7 priming site/V5 epitope) were used. The sizes of the PCR products were checked by electrophoresis on 1% agarose gel.

### *In vitro* CEST-MRI of cell lysates

The rdLRP plasmid, under control of the cytomegalovirus (CMV) promoter, was transfected into HEK293T and GL261N4 cells using lipofectamine. After 24 hours incubation, the cells were lysed in pH 7.5 RIPA buffer and underwent 3 freeze/thaw/ sonication cycles. Protein content of the supernatant was measured and protein concentrations of control and LRP transfected cell lysates were adjusted to be equivalent (typically 10 *µ*g/*µ*l protein). Control and rdLRP cell lysate samples were loaded into 5 mm od sample tubes and placed into 50 ml Falcon tubes containing water. CEST Z-spectra were acquired on a 4.7T MRI scanner (Bruker Biospin, Ettlingen, Germany) using a CEST-EPI pulse sequence, with flip-angle = 90° and TE/TR = 20/8000 ms. The saturation pulse power, duration, and frequency offsets were 3.6 *µ*T, 5000 ms, and 7 to −7 ppm with 0.25 ppm increments, respectively.

### Animal model

Six-to-8-week-old female C57/BL6 mice were purchased from Envigo (n=10). All animal experiments were performed in accordance with protocols approved by the Brigham and Women’s Hospital Institutional Animal Care and Use Committee (IACUC). 100,000 cells of GL261N4 / rdLRP-GL261N4 were implanted stereotactically with a Kopf stereotactic frame (David Kopf Instruments) in the right frontal lobe of the mice (ventral 3.0 mm, rostral 0.5 mm, and right lateral 2.0 mm from bregma).

### *In vivo* CEST MRI

The rdLRP was expressed in a murine glioma cell line and transplanted into mice brains as described above (n=10 mice, 50% served as control). Imaging was performed at 11 and 18 days post implantation using a 7T preclinical MRI (Bruker Biospin, Ettlingen, Germany). The mice were anesthetized using 0.5 to 2% Isoflurane during the imaging on a dedicated heated cradle, and the respiration rate was continuously monitored using a mechanical sensor. T_2_-weighted images were acquired using a multi-slice multi-echo (MSME) protocol, with TR/TE = 60/2000 ms. T_2_ maps were acquired using the multi-echo spin-echo protocol, TR = 2000 ms, 25 TE values between 8-200 ms. Z-spectra were acquired using the same CEST-EPI sequence used for the cell lysate *in vitro* study, with a saturation pulse power of 0.7 *µ*T. Static magnetic field B_0_ maps were acquired using the WASSR method^43^, employing a saturation pulse power of 0.3 *µ*T, TR/TE = 8000/20 ms, flip-angle = 90°, saturation duration = 3000 ms, and a saturation pulse frequency varying between −1 to 1 ppm in 0.1 ppm increments. The field of view (19 mm × 19 mm × 1 mm) and image resolution (297 × 297 × 1000 *µ*m^3^) were identical for all scans.

### Statistical analysis

Group analysis was carried out using a two-tailed student t-test, calculated using the open source SciPy scientific computing library for Python^44^. Differences were considered significant at P<0.05. Box plots (Fig. 4c,h) represent the median, interquartile range, Tukey style whiskers, outliers (yellow circles), and individual data points.

### Image analysis

Data analysis was performed using MATLAB R2018a and Python 3.6 custom written scripts, based on previously published routines, as described below. T_2_ maps reconstruction was performed using exponential fitting. CEST images were corrected for B_0_ inhomogeneity using the WASSR method^43, 45^, followed by cubic spline smoothing^46^. The magnetization transfer ratio asymmetry curve was calculated using: MTR_*asym*_ = (S^− Δ *ω*^ – S^+Δ *ω*^) / S_0_, where S^±Δ *ω*^ is the signal measured with saturation offset of ±Δ *ω* around the water frequency and S_0_ is the unsaturated signal. Semi-quantitative mapping of the CEST molecular compound amplitudes was performed using a 5-pool (water, semi-solid macro-molecules, amide, rNOE, and amine) Lorentzian fitting model, with the starting point and boundaries described in^47^. For amide proton amplitude image visualization, an image down-sampling expedited adaptive least-squares (IDEAL) fitting approach was implemented, as described in^48^. Mouse tumor regions of interest (ROIs) were manually delineated based on the T_2_ weighted images. Finally, the amide Lorentzian peak amplitude pixel values were overlaid atop the T_2_ weighted images at the tumor ROIs (Fig. 4).

### Data availability

The datasets generated and analysed during the current study are available from the corresponding author upon request.

### Code Availability

Source code is available from the corresponding author upon request.

---

**Extended Data Fig. 1.**
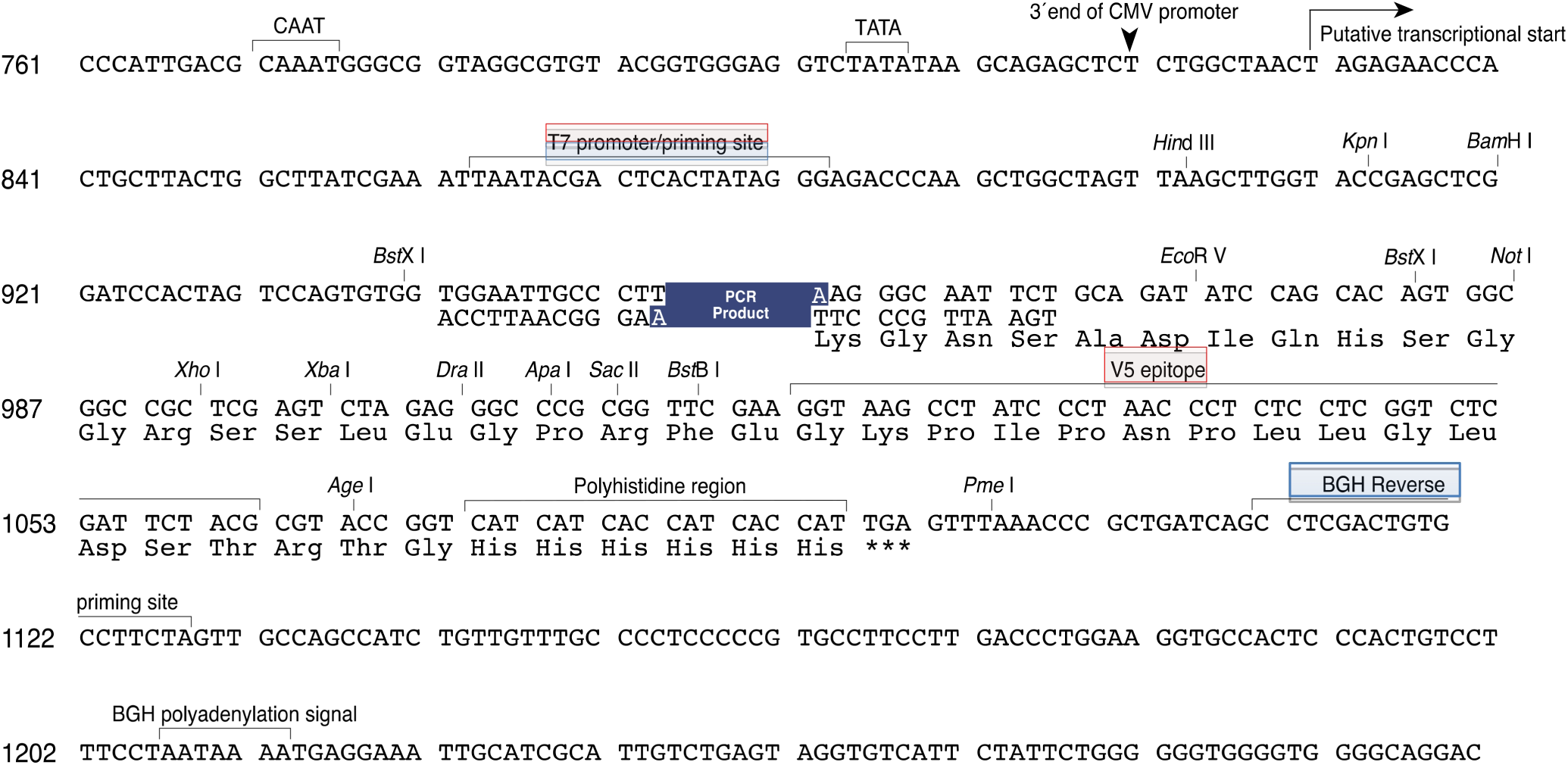
rdLRP DNA sequence.

**Extended Data Fig. 2.**
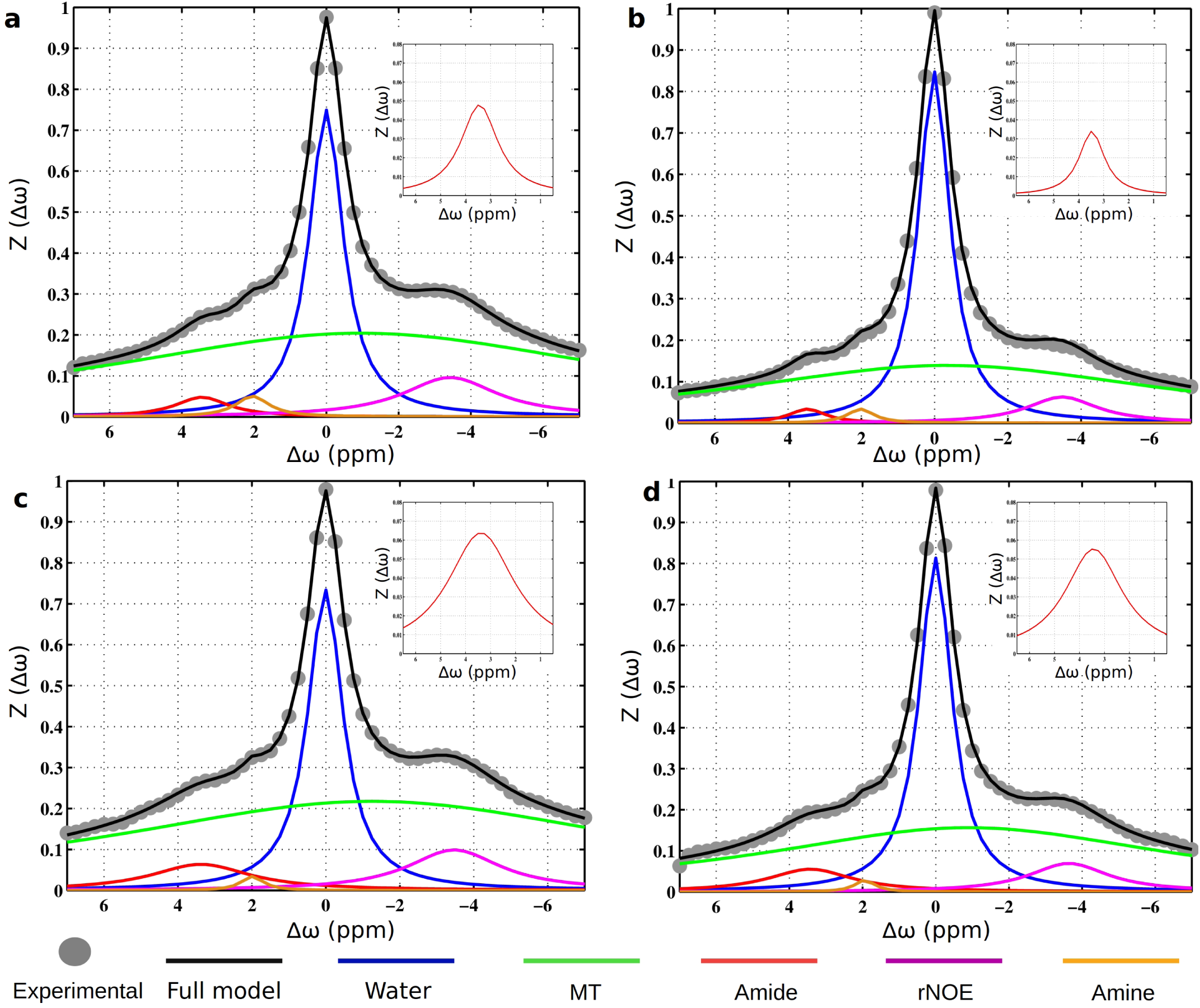
Extraction of the CEST amide proton amplitude *in vivo* using a Lorentzian fitting model. Each sub-figure represent the fitting of the experimentally measured z-spectra in the contralateral (left, **a**,**c**) and tumor (right, **b**,**d**) to a 5 pool model (water, semi-solid macro-molecules (MT), amide, aliphatic (rNOE), and amine), for the representative control (top row, **a-b**) and rdLRP (bottom row, **c-d**) mice, presented in Fig. 4. The insets provide a zoom-in display of the amide signal contribution.

